## Supplementary material for "Quantitative insights into the cyanobacterial cell economy": All supplementary and source data files: Zavrel2018_Fig1_Supplement_1_Nutrients_sufficiency_R1.pdf

| Ratio of the requirements of selected elements by *Synechocystis* cells and the elements supplemented by the replacement of a spare culture medium during turbidostat experiments in this study. | | | | | | | | |
|---|---|---|---|---|---|---|---|---|
| **Red light intensity [μE $m^{-2}$ $s^{-1}$]** | **27.5** | **55** | **110** | **220** | **440** | **660** | **880** | **1100** |
| **Na** | 0.2% | 0.4% | 0.4% | 0.5% | 0.4% | 0.4% | 0.5% | 0.5% |
| **N** | 6.1% | 9.6% | 10.8% | 12.9% | 9.7% | 11.6% | 14.4% | 12.7% |
| **Mg** | 16.8% | 26.4% | 29.7% | 35.6% | 26.7% | 31.9% | 39.7% | 35.1% |
| **S** | 9.4% | 14.7% | 16.6% | 19.9% | 14.9% | 17.8% | 22.2% | 19.6% |
| **Ca** | 20.4% | 31.9% | 36.0% | 43.1% | 32.4% | 38.6% | 48.1% | 42.5% |
| **Fe** | 40.3% | 63.1% | 71.2% | 85.1% | 64.0% | 76.4% | 95.1% | 83.9% |
| **P** | 33.3% | 52.2% | 58.9% | 70.5% | 52.9% | 63.2% | 78.7% | 69.5% |

| Estimated uptake rates of selected elements by *Synechocystis* cells during the turbidostat cultivation [mg $L^{-1}$ $h^{-1}$]. The calcualtions are based on direct measurements of cellular dry weights and specific growth rates in this study, and on maximal concentration of particular elements in *Synechocystis* biomass as recorded in the literature. | | | | | | | | |
|---|---|---|---|---|---|---|---|---|
| **Red light intensity [μE $m^{-2}$ $s^{-1}$]** | **27.5** | **55** | **110** | **220** | **440** | **660** | **880** | **1100** |
| **Na** | 0.03 | 0.05 | 0.08 | 0.14 | 0.18 | 0.21 | 0.20 | 0.17 |
| **N** | 0.42 | 0.80 | 1.27 | 2.23 | 2.89 | 3.37 | 3.18 | 2.69 |
| **Mg** | 0.03 | 0.07 | 0.10 | 0.18 | 0.24 | 0.28 | 0.26 | 0.22 |
| **S** | 0.03 | 0.05 | 0.08 | 0.14 | 0.18 | 0.21 | 0.20 | 0.17 |
| **Ca** | 0.06 | 0.11 | 0.17 | 0.30 | 0.38 | 0.45 | 0.42 | 0.36 |
| **Fe** | 0.01 | 0.03 | 0.04 | 0.07 | 0.09 | 0.11 | 0.10 | 0.09 |
| **P** | 0.05 | 0.10 | 0.15 | 0.27 | 0.35 | 0.40 | 0.38 | 0.32 |

| Refilling rates of selected elements during *Synechocystis* cultivation in the turbidostat regime, based on data from this study [mg $L^{-1}$ $h^{-1}$]. | | | | | | | | |
|---|---|---|---|---|---|---|---|---|
| **Red light intensity [μE $m^{-2}$ $s^{-1}$]** | **27.5** | **55** | **110** | **220** | **440** | **660** | **880** | **1100** |
| **Na** | 11.52 | 14.08 | 19.71 | 28.91 | 49.90 | 48.78 | 36.99 | 35.45 |
| **N** | 6.88 | 8.41 | 11.77 | 17.26 | 29.80 | 29.13 | 22.09 | 21.17 |
| **Mg** | 0.21 | 0.25 | 0.35 | 0.52 | 0.89 | 0.87 | 0.66 | 0.63 |
| **S** | 0.28 | 0.34 | 0.48 | 0.70 | 1.21 | 1.18 | 0.89 | 0.86 |
| **Ca** | 0.27 | 0.33 | 0.47 | 0.69 | 1.18 | 1.16 | 0.88 | 0.84 |
| **Fe** | 0.03 | 0.04 | 0.06 | 0.08 | 0.14 | 0.14 | 0.11 | 0.10 |
| **P** | 0.15 | 0.18 | 0.26 | 0.38 | 0.65 | 0.64 | 0.49 | 0.47 |

| Weights of selected elements in *Synechocystis* cells, based on directly measured cellular dry weight in this study, and on maximal concentration of particular elements in *Synechocystis* biomass as recorded in the literature [mg $L^{-1}$ $h^{-1}$]. | | | | | | | | |
|---|---|---|---|---|---|---|---|---|
| **Red light intensity [μE $m^{-2}$ $s^{-1}$]** | **27.5** | **55** | **110** | **220** | **440** | **660** | **880** | **1100** |
| **Na** | 1.04 | 1.31 | 1.37 | 1.73 | 1.75 | 2.05 | 2.03 | 1.82 |
| **N** | 16.5 | 20.7 | 21.6 | 27.4 | 27.7 | 32.5 | 32.0 | 28.9 |
| **Mg** | 1.36 | 1.71 | 1.78 | 2.26 | 2.28 | 2.68 | 2.64 | 2.38 |
| **S** | 1.03 | 1.29 | 1.35 | 1.71 | 1.73 | 2.03 | 2.00 | 1.80 |
| **Ca** | 2.19 | 2.75 | 2.87 | 3.64 | 3.67 | 4.31 | 4.26 | 3.83 |
| **Fe** | 0.53 | 0.66 | 0.69 | 0.88 | 0.89 | 1.04 | 1.03 | 0.92 |
| **P** | 1.98 | 2.48 | 2.59 | 3.29 | 3.32 | 3.90 | 3.85 | 3.46 |

| Parameters of *Synechocystis* cultures as measured during the turbidostat experiments in this study. | | | | | | | | |
|---|---|---|---|---|---|---|---|---|
| **Red light intensity used in this study [μE $m^{-2}$ $s^{-1}$]** | **27.5** | **55** | **110** | **220** | **440** | **660** | **880** | **1100** |
| **Specific growth rate μ measured in this study [$h^{-1}$]** | 0.025 | 0.039 | 0.059 | 0.081 | 0.104 | 0.104 | 0.099 | 0.093 |
| **Flow rate of spare cultivation media measured in this study [$h^{-1}$]** | 0.028 | 0.034 | 0.048 | 0.070 | 0.121 | 0.118 | 0.089 | 0.086 |
| **Dry weight of *Synecocystis* cells measured in this study [mg $L^{-1}$]** | 132 | 166 | 173 | 220 | 221 | 260 | 256 | 231 |

| Elemental composition of *Synechocystis* cells based on data available in the literature. | | | | | |
|---|---|---|---|---|---|
| **Element** | **BG11 [mg $L^{-1}$]** | **Content in cell [% of DW] Min** | **Content in cell [% of DW] Max** | **Maximal value recorded in the literature** | **Reference** |
| **Na** | 413.4 | 0.1 | 0.8 | 0.8 | Touloupakis et al. 2016 |
| **N** | 246.9 | 8.0 | 11.2 | 12.5 | Zavřel et al. 2017 |
| | | 10.0 | 11.1 | | Touloupakis et al. 2015 |
| | | 10.2 | 11.5 | | Touloupakis et al. 2016 |
| | | 11.3 | 11.3 | | Shastri and Morgan 2005 |
| | | 12.5 | | | Kim et al. 2011 |
| | | 7.1 | 7.7 | | Blom 2014 |
| **S** | 10.0 | 0.4 | 0.8 | 0.8 | Zavřel et al. 2017 |
| | | 0.4 | 0.4 | | Touloupakis et al. 2015 |
| | | 0.4 | 0.4 | | Touloupakis et al. 2016 |
| | | 0.7 | | | Kim et al. 2011 |
| **Ca** | 9.8 | 0.3 | 1.7 | 1.7 | Touloupakis et al. 2016 |
| **Mg** | 7.4 | 0.4 | 1.0 | 1.0 | Touloupakis et al. 2016 |
| **P** | 5.4 | 1.5 | | 1.5 | Kim et al. 2011 |
| **Fe** | 1.2 | 0.1 | 0.4 | 0.4 | Cheng and He 2014 |
| | | 0.4 | | | Kim et al. 2011 |
