## Supplementary material for "Quantitative insights into the cyanobacterial cell economy": All supplementary and source data files: Zavrel2018_Fig5_Supplement_1_model summary.pdf

### SMALL–SCALE PROTEOME ALLOCATION MODEL FOR PHOTOTROPHIC GROWTH

| parameter | definition | value | source |
|---|---|---|---|
| $P_m$ | cell membrane permeability to inorganic carbon | 0.108 [ $dm\ h^{-1}$ ] | (2) |
| $A_{cell}$ | cell surface area | $1.26 \cdot 10^{-9}$ [ $dm^2\ cell^{-1}$ ] | This study |
| $V_{cell}$ | cell volume | $4.19 \cdot 10^{-15}$ [ $dm^3\ cell^{-1}$ ] | This study |
| $N_A$ | Avogadro constant | $6.022 \cdot 10^{23}$ [ $mol^{-1}$ ] | |
| $k^t_{cat}$ | maximal import rate | 43560 [ $h^{-1}$ ] | (3) |
| $K_t$ | half-saturation constant of the transporter enzyme | 15 [ µM ] | (4) |
| $k^m_{cat}$ | maximal metabolic rate | 32700 [ $h^{-1}$ ] | (5) |
| $K_m$ | half-saturation constant of the metabolic enzyme | 2441560 [ molecules $cell^{-1}$ ] | (5) |
| $\gamma_{max}$ | maximal translation rate | 79200 [ aa $h^{-1}$ $molecules^{-1}$ ] | (6) |
| $K_a$, $K_e$ | half-saturation constant of amino acids and energy units for each reaction | 10000 [ molecules $cell^{-1}$ ] | (1) |
| $d_p$ | protein half-life | 1/23 [ $h^{-1}$ ] | (7) |
| σ | effective absorption cross-section of the photosynthetic unit | 0.7 [ $nm^2$ ] | This study |
| τ | maximal turnover rate of the photosynthetic unit | 270000 [ $h^{-1}$ ] | This study |
| $k_d$ | rate constant for photodamdage | $10^{-6}$ | This study |
| $m_v$ | energy maintenance rate | $7 \cdot 10^9$ [ molecules $cell^{-1}$ $h^{-1}$ ] | (8) |
| $D_c$ | average cell density (protein mass per cell) | $1.4 \cdot 10^{10}$ [ aa $cell^{-1}$ ] | (1) |
| $n_R$ | ribosome length | 7358 [ aa $molecule^{-1}$ ] | (1) |
| $n_Q$ | average protein length for house-keeping proteins | 300 [ aa $molecule^{-1}$ ] | This study |
| $n_P$ | length of one photosynthetic unit | 95451 [ aa $molecule^{-1}$ ] | (1) |
| $n_T$ | transporter length | 1681 [ aa $molecule^{-1}$ ] | (1) |
| $n_M$ | length of one metabolic enzyme complex | 28630 [ aa $molecule^{-1}$ ] | (1) |
| $m_\alpha$ | amount of energy units consumed to create one amino acid | 45 | (1) |
| $m_c$ | average carbon chain length of an amino acid | 5 | (1) |
| $m_\gamma$ | amount of energy units needed for one translational elongation step | 3 | (1) |
| $m_\Phi$ | amount of energy units produced during photosynthesis | 8 | (1) |

*Proteome Allocation Problem*

$$\max_{\beta,\ X,\ \mu} \quad \mu$$

$$s.t.\ \frac{d[X]}{dt} - \mu \cdot X = 0 ,$$

$$\sum_j \beta_j = 1 ,\ \ \forall j \in \mathbb{E} :\ \beta_j \geq 0 ,$$

$$\sum_j n_j \cdot [j] + [aa] + \frac{[c_i]}{m_c} = D_c ,$$

$$n_Q \cdot [Q] = 0.5 \cdot D_c ,$$

$$\mathbb{E} = \{R, Q, P, T, M\} ,$$

$$X = [ci, aa, e, Q, P^o, P*, T, M, R]^T \in \mathbb{R}_+ .$$

*ODE System*

$$\frac{d[c_i]}{dt} = v_d + v_t - m_c \cdot v_m ,$$

$$\frac{d[aa]}{dt} = v_m + n_P \cdot v_i - \sum_j n_j \cdot \gamma_j + d_p \cdot \sum_j n_j \cdot [j] ,$$

$$\frac{d[z]}{dt} = \gamma_z - d_P \cdot [z] ,$$

$$\frac{d[P^o]}{dt} = \gamma_P - v_1 + v_2 - d_p \cdot [P^o] ,$$

$$\frac{d[P*]}{dt} = v_1 - v_2 - v_i - d_p \cdot [P*] ,$$

$$\frac{d[e]}{dt} = m_\phi \cdot v_2 - v_t - m_\mu \cdot v_m - m_\gamma \cdot \sum_j n_j \cdot \gamma_j - \frac{m_v \cdot [e]}{10 + [e]} ,$$

$$\forall j \in \mathbb{E} , \forall z \in \mathbb{E} \backslash P .$$

*Reaction Rates*

$$v_d = P_m \cdot \frac{A_{cell}}{V_{cell}} \cdot (N_A \cdot V_{cell} \cdot [c_i^x] - [c_i]) ,$$

$$v_t = [T] \cdot k^t_{cat} \cdot \frac{[c_i^x]}{K_t + [c_i^x]} \cdot \frac{[e]}{K_e + [e]} ,$$

$$v_m = [M] \cdot k^m_{cat} \cdot \frac{[c_i]}{K_m + [c_i]} \cdot \frac{[e]}{K_e + [e]} ,$$

$$\gamma_j = [R] \cdot \beta_j \cdot \frac{\gamma_{max}}{n_j} \cdot \frac{[aa]}{K_a + [aa]} \cdot \frac{[e]}{K_e + [e]} ,$$

$$v_1 = \sigma \cdot light \cdot [P^o] ,$$

$$v_2 = \tau \cdot [P*] ,$$

$$v_i = k_d \cdot \sigma \cdot light \cdot [P*] ,$$

$$\forall j \in \mathbb{E} .$$
