## Supplementary material for "Quantitative insights into the cyanobacterial cell economy": All supplementary and source data files: Zavrel2018_Fig6_Supplement_1_WB_antibodies_blots.pdf

Table 1. List of antibodies used in this study.

| Antibody | Agrisera catalogue number | Dilution | Protein apparent MW |
|---|---|---|---|
| Rabbit Anti-RbcL (Rubisco large subunit, form I and form II) | AS03 037 | 1:5000 | 52.5 kDa |
| Rabbit Anti-PsaC (PSI-C core subunit of photosystem I) | AS10 939 | 1:1000 | 9 kDa |
| Rabbit Anti-PsbA (D1 protein of PSII, C-terminal) | AS05 084 | 1:10000 | 28-30 kDa |
| Rabbit Anti-S1 (30S ribosomal protein S1) | AS08 309 | 1:2000 | 35 kDa |
| Rabbit Anti-L1 (50S ribosomal protein L1) | AS11 1738 | 1:1000 | 25 kDa |

Table 2. List of protein standards used in this study.

| Standard | Agrisera catalogue number | Protein apparent MW | Concentrations |
|---|---|---|---|
| Purified spinach RbcL | AS01 017S | 52.7 kDa | 0.375, 0.75, 1.5 pmol |
| Recombinant PsaC from *Synechocystis* PCC 6803 | AS04 042S | 11.5 kDa | 0.075, 0.3, 0.6 pmol |
| Recombinant PsbA from *Synechocystis* PCC 6803 | AS01 016S | 41.5 kDa | 0.125, 0.5, 1 pmol |

### PsbA: PSII

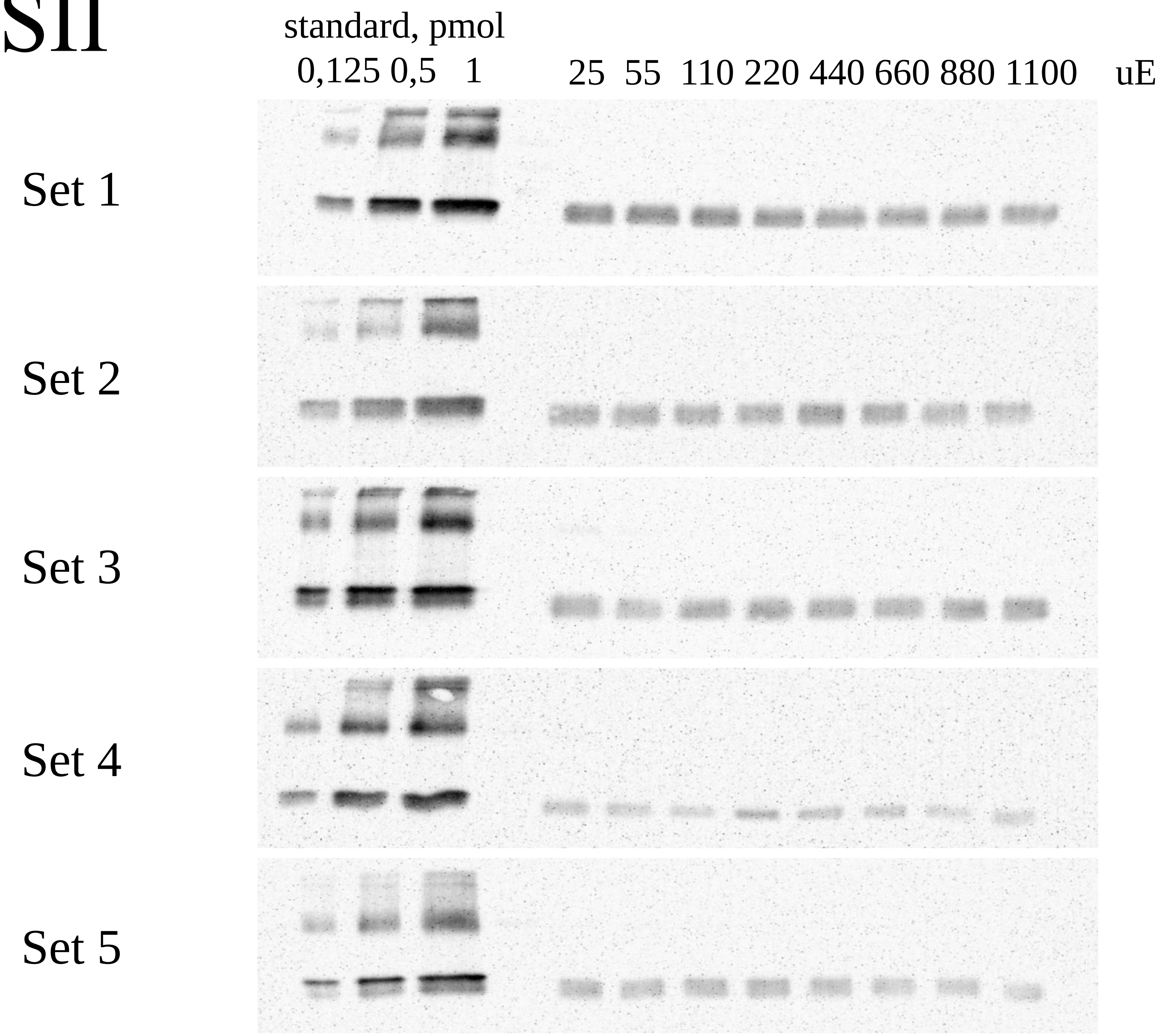
standard, pmol
0,125 0,5 1
25 55 110 220 440 660 880 1100 uE
Set 1
Set 2
Set 3
Set 4
Set 5

### PsaC: PS I

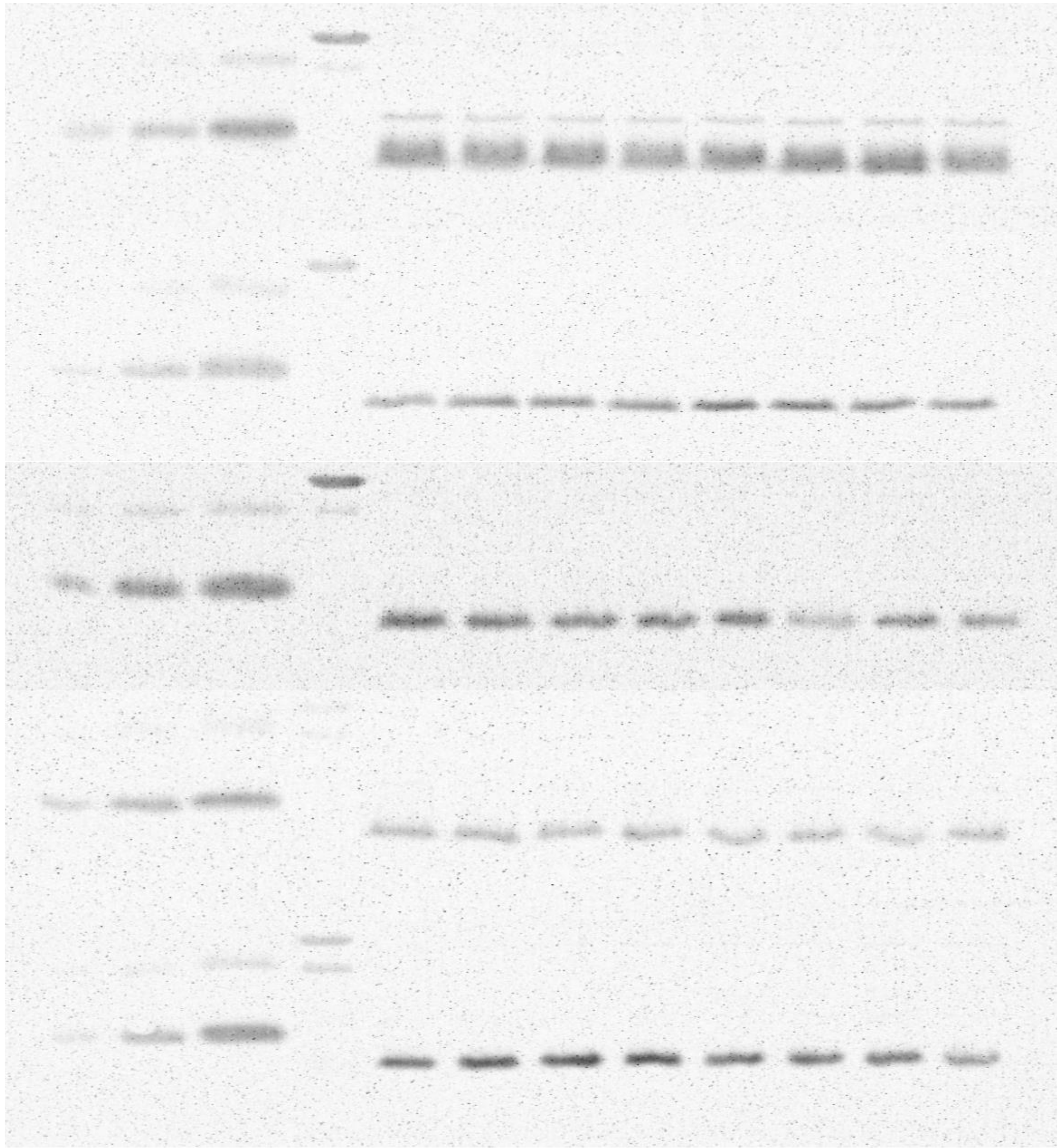
standard, pmol
0,075 0,3 0,6
25 55 110 220 440 660 880 1100 uE
Set 1
Set 2
Set 3
Set 4
Set 5

### RbcL: RuBisCO

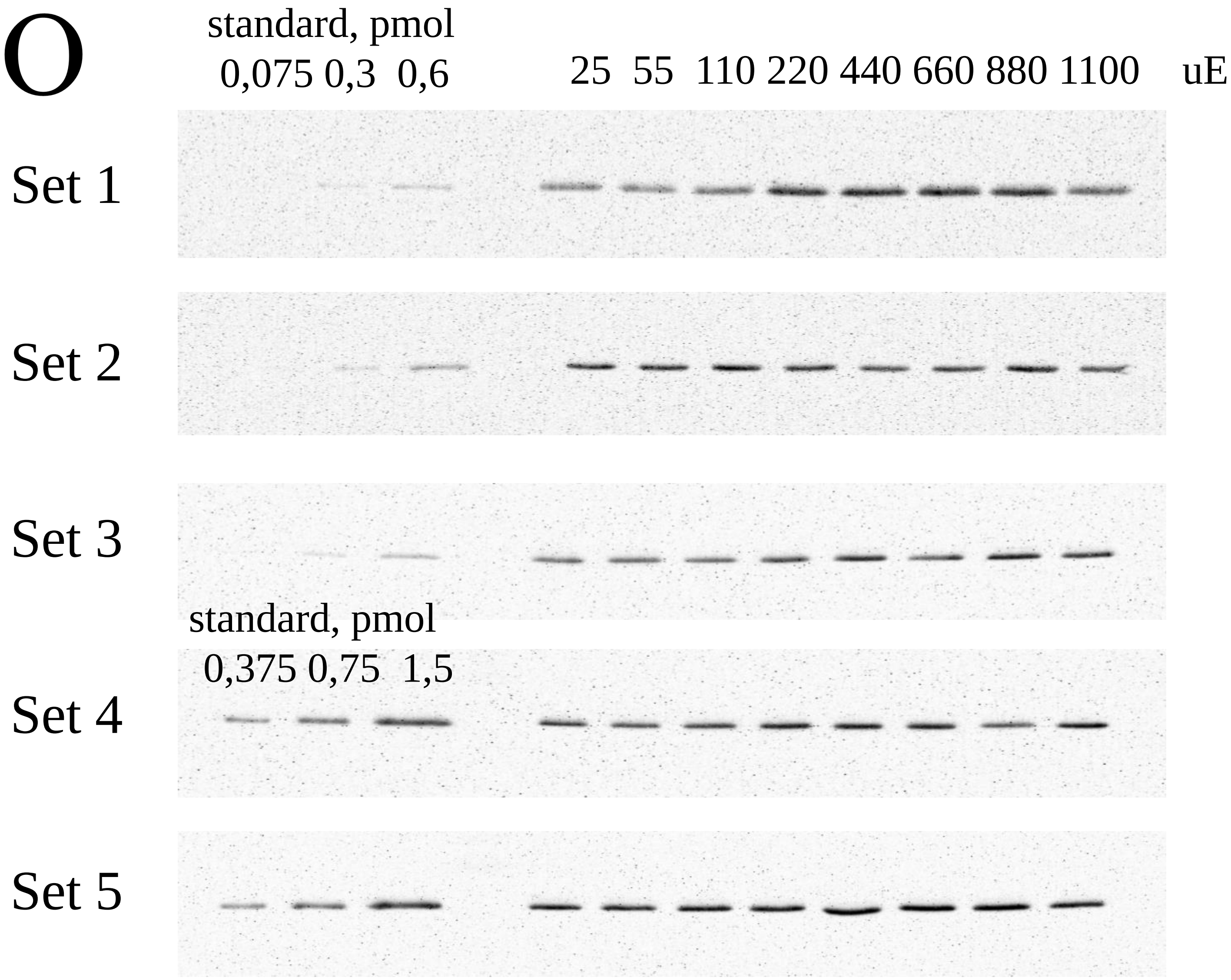

### L1: ribosome large subunit

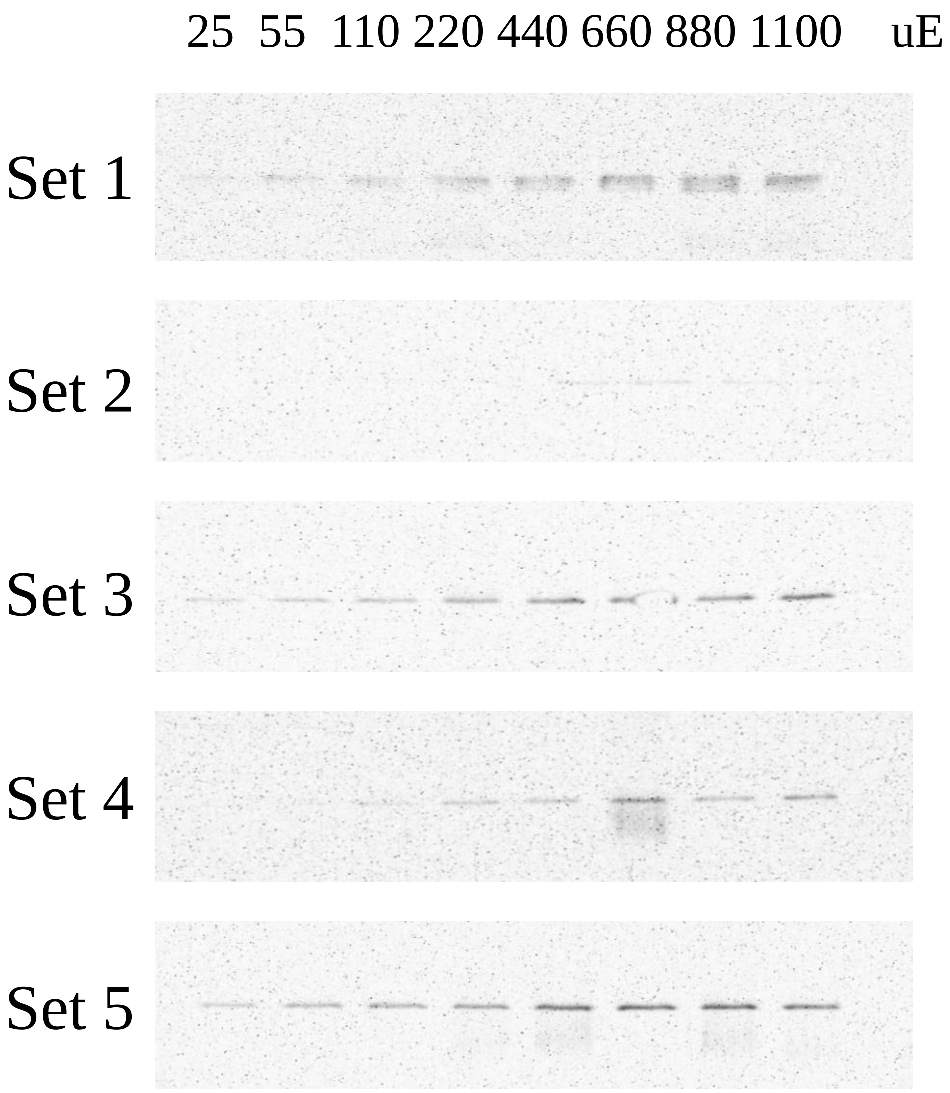

### S1: ribosome small subunit

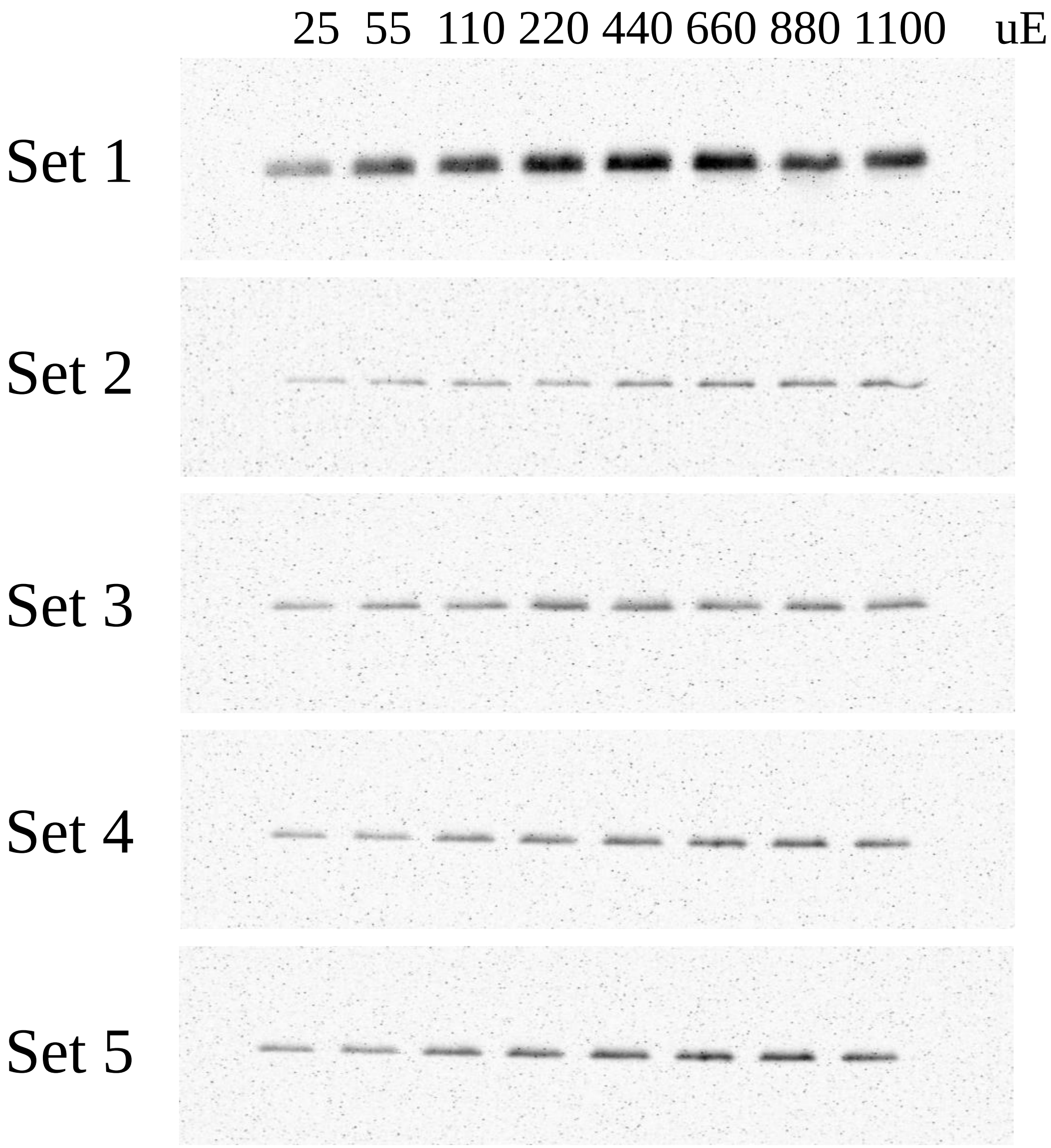
