## Supplementary figures and images for "Quantitative insights into the cyanobacterial cell economy"

### Zavrel2018_Fig2_Supplement_1_Data_per_light_intensity_R1.pdf

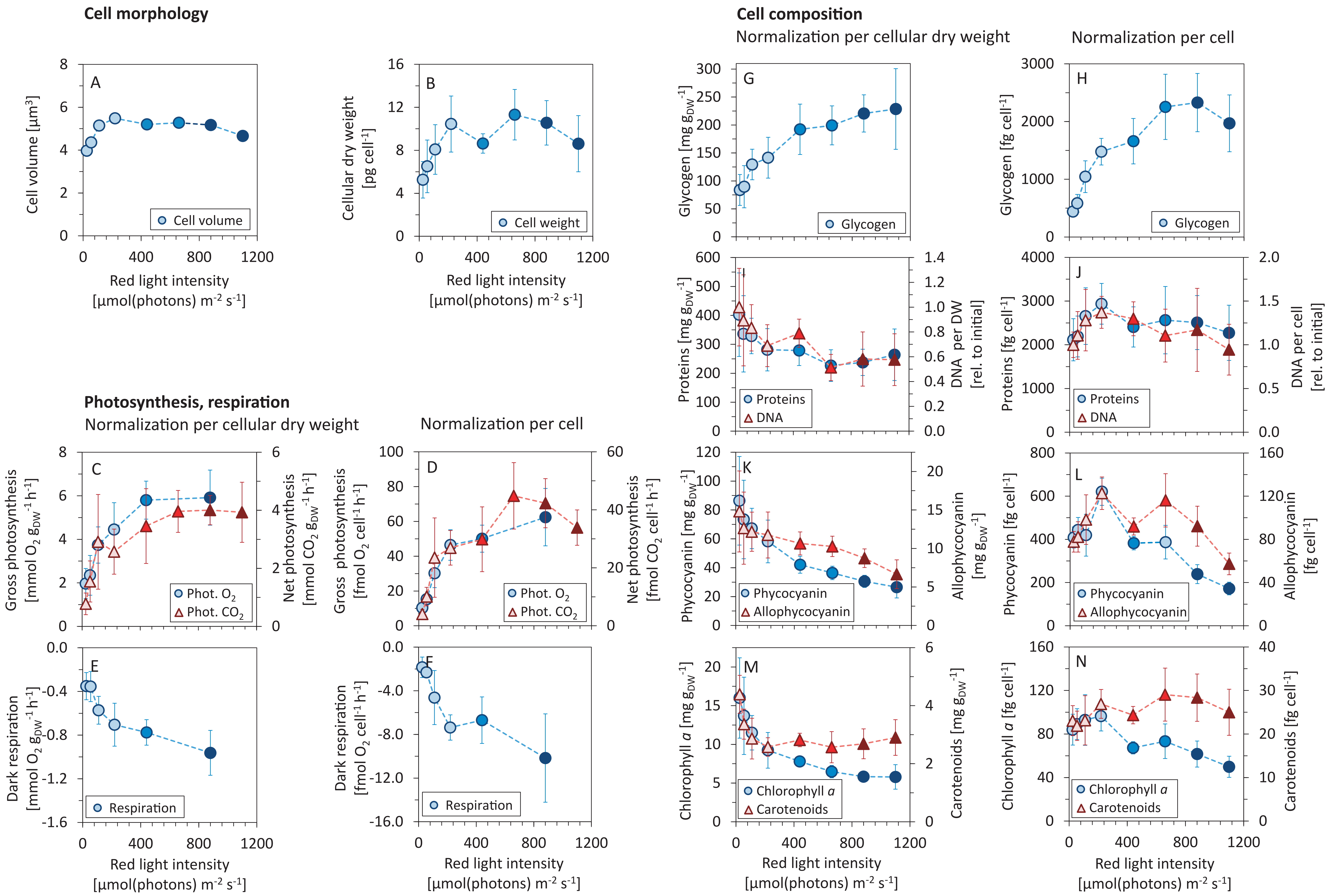

### Zavrel2018_Fig3_Supplement_3_Elbow_method_find_k7.pdf

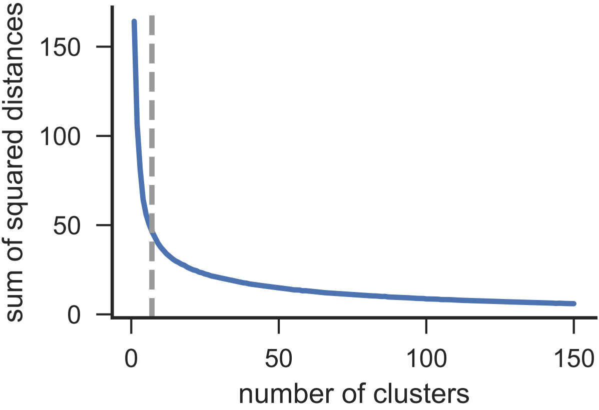

### Zavrel2018_Fig4_Supplement_1_proteomaps_levels_2_3_4.pdf

**Processes** (Proteomaps level 2)

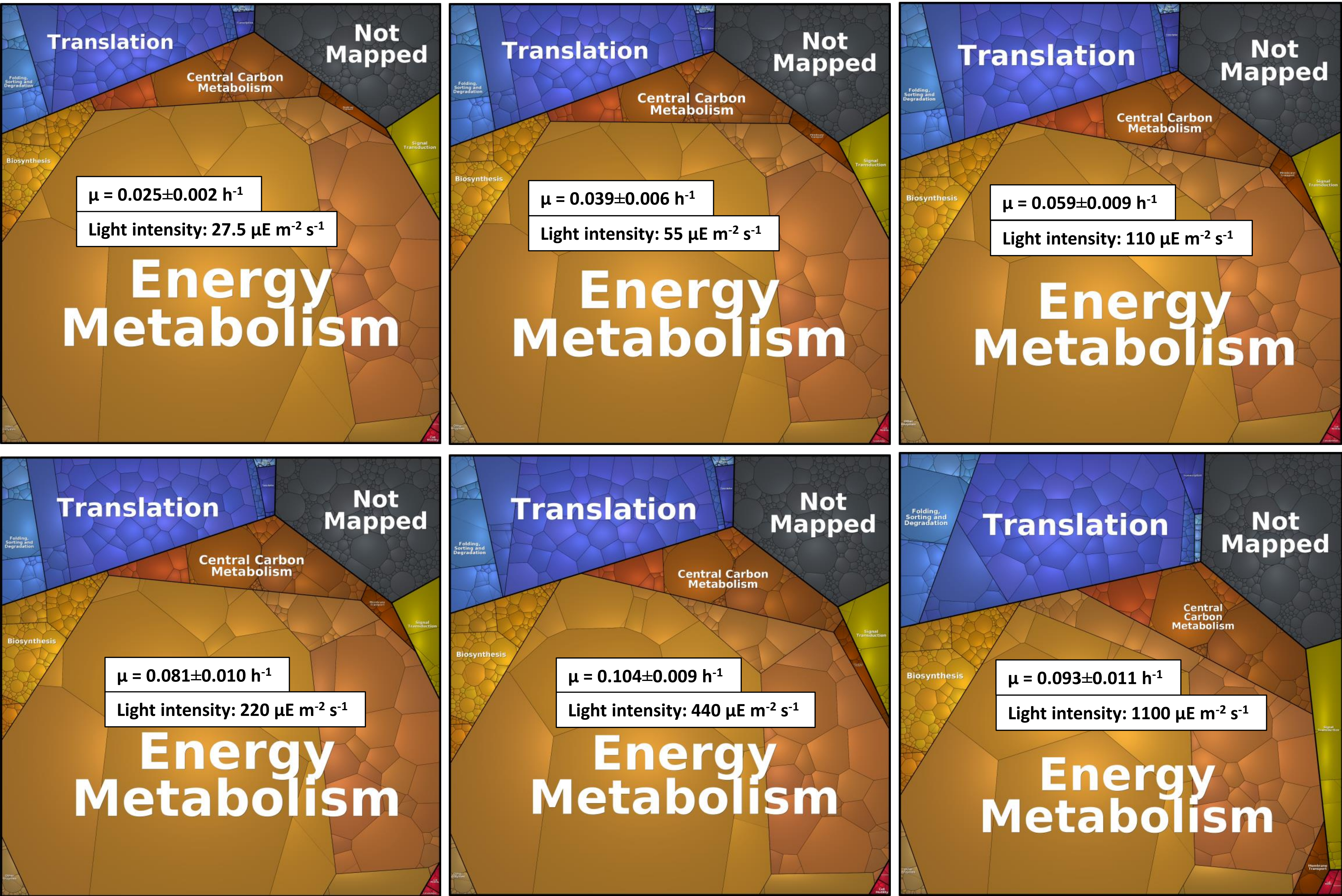

**Processes** (Proteomaps level 3)

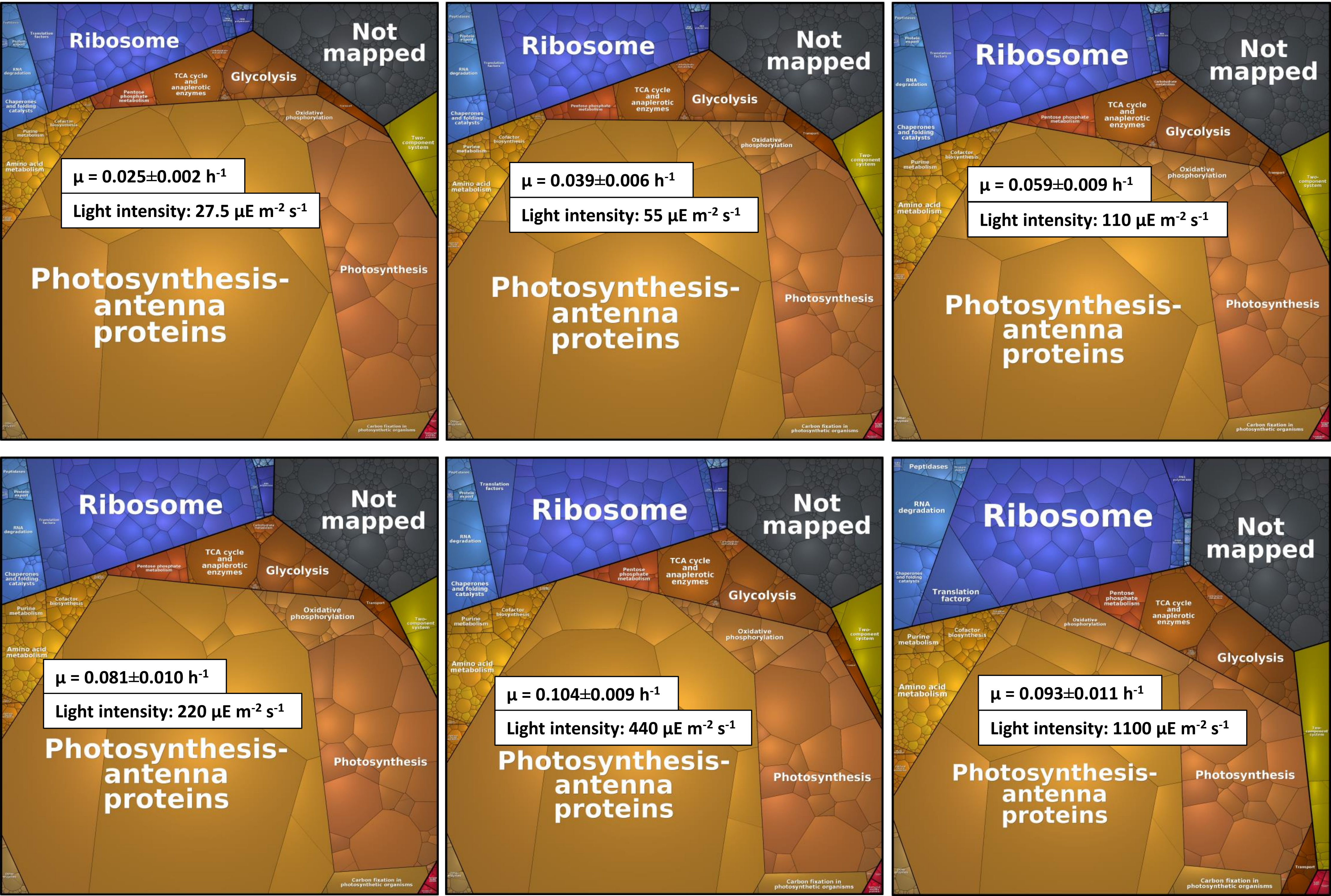

**Proteins** (Proteomaps level 4)

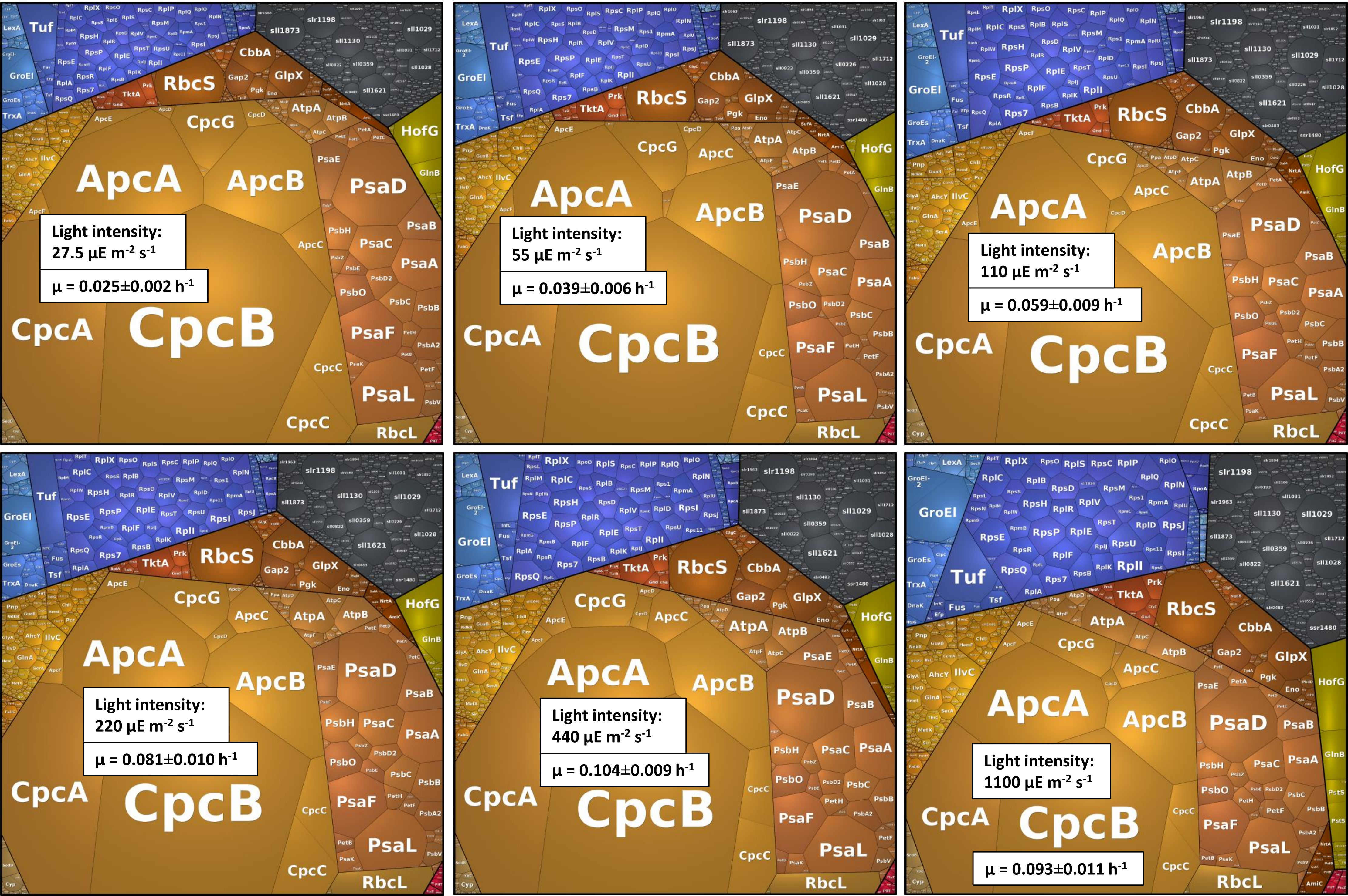

### Zavrel2018_Fig6_Supplement_2_Constant proteome fractions.pdf

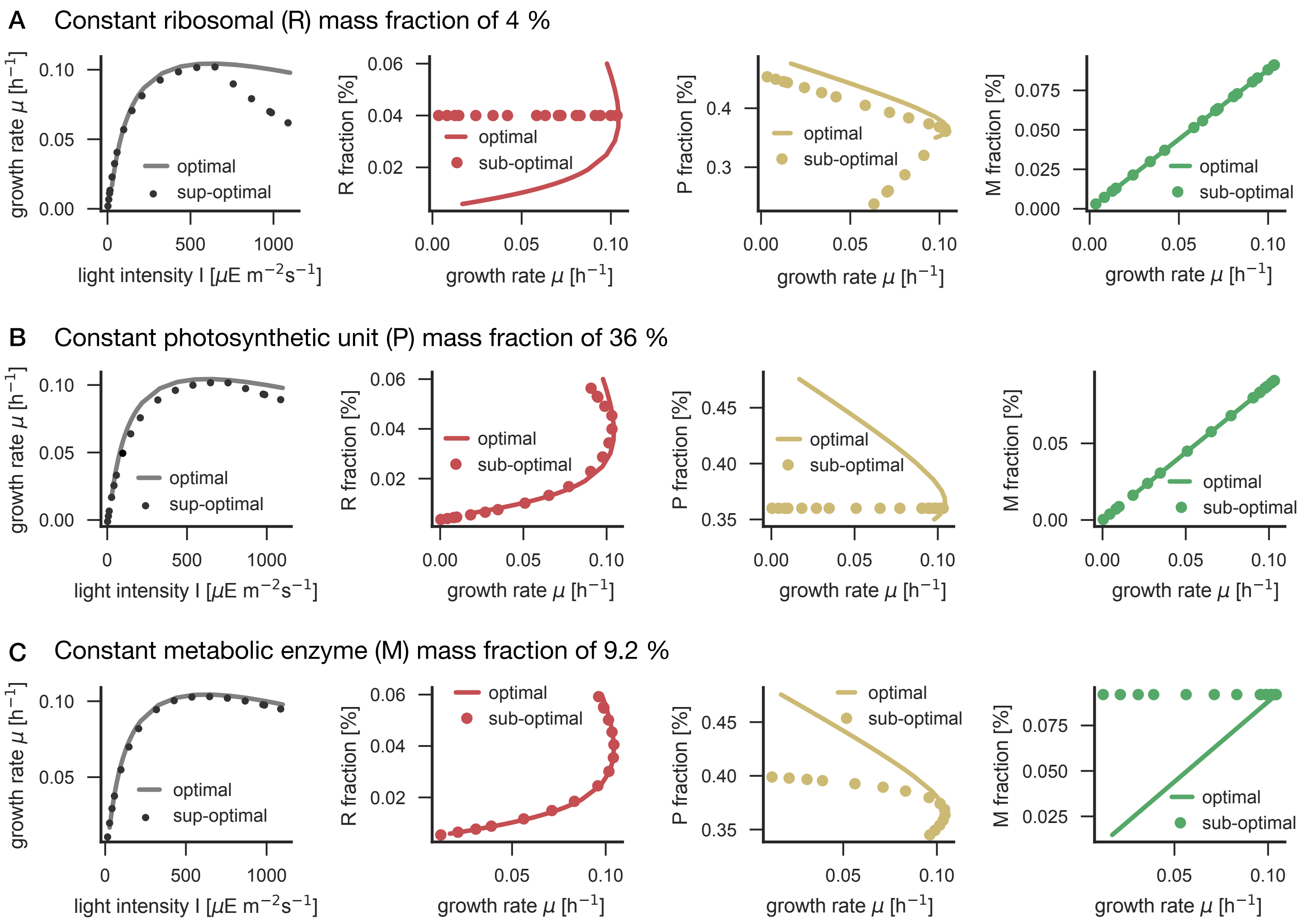
